# Optimizing DNA extraction from environmentally degraded bone samples for molecular identification of cetacean species

**DOI:** 10.64898/2026.08.31.748265

**Authors:** Daniel Pagani, Sara M. Rodríguez, Maribet Gamboa

## Abstract

Molecular identification of cetacean bone remains can be limited by DNA degradation and the presence of PCR inhibitors. Here, we present an optimized DNA extraction protocol based on a total demineralization method for environmentally exposed cetacean bones. The protocol uses 100 mg of bone powder, 24 h digestion with EDTA, N-lauroylsarcosine, and proteinase K, followed by a modified silica-column purification. Nine environmentally degraded bone samples representing eight individuals were processed. DNA concentrations ranged from 7.3 to 57.1 ng/µL (mean ± SD = 25.91 ± 13.91 ng/µL). The mitochondrial cytochrome b gene was successfully amplified from all samples using conventional PCR, and five samples (55.6%) yielded sequences suitable for downstream analysis. BLASTn identified *Balaenoptera physalus* as the closest database match for all recovered sequences, and phylogenetic analysis further supported their association with *B. physalus* reference sequences. These results demonstrate that the proposed protocol provides a practical approach for recovering amplifiable and molecularly informative mitochondrial DNA from environmentally degraded cetacean bone material, facilitating molecular identification from challenging skeletal remains.

## Introduction

Molecular identification of cetacean bone remains is relevant to ecological, forensic and conservation studies, particularly when morphological identification is limited by fragmentation, environmental exposure or poor preservation of cetacean bones (Speller et al. 2016). Molecular approaches can overcome some of these limitations, however, DNA recovered from bone tissue is often fragmented, present at low copy number and affected by co-extracted inhibitory compounds such as calcium and humic acids, which can compromise the PCR amplification (Kalmár et al. 2000; Ambers et al. 2014).

Bone DNA extraction therefore requires efficient disruption of the mineral matrix while minimizing DNA loss and the carry-over of inhibitors. In this context, commercial kits based on silica columns provide a simple and accessible purification strategy, although their performance may be reduced when applied to degraded material because they can co-purify PCR inhibitors with the DNA, requiring larger sample volumes and multiple wash steps, which may increase the risk of DNA loss (Duijs and Sijen 2020). Total demineralization, as described by Loreille et al. (2007), was developed to improve DNA recovery from human skeletal remains and has subsequently been evaluated in forensic and ancient DNA workflows (Xavier et al. 2021).

In this study, we present an optimized DNA extraction protocol for environmentally exposed cetacean bones based on the total demineralization approach of Loreille et al. (2007). We adapted the digestion conditions and modified the E.Z.N.A Tissue DNA Kit protocol to recover DNA from cetacean bones. Protocol performance was evaluated based on DNA concentration and purity, conventional PCR amplification of the mitochondrial cytochrome b gene, sequencing success and downstream molecular identification.

## Methods

### Bone samples

Bone remains were obtained by the Servicio Nacional de Pesca y Acuicultura (SERNAPESCA), and Huachipato steel plant in Biobio region, Chile. DNA extraction and molecular identification were performed on a subset of nine cetacean bone samples, including skull, mandible and vertebrae. These nine samples perhaps are represented in eight individuals, because the skull and vertebra from Huachipato were known to originate from the same individual. All specimens correspond to cetacean bone remains found through bycatch by bottom trawling nets and strandings along the coast. All samples lacked prior taxonomic identification. Bone powder was obtained by perforating a portion of each bone fragment using a dental drill, which was cleaned with 95 % ethanol before each use. The resulting bone powder and fragments were collected and transferred to sterile 1.5 mL Eppendorf tubes until DNA extraction.

### DNA extraction and protocol optimization

The DNA extraction was based on the total demineralization protocol described by Loreille et al. (2007), originally developed for human skeletal remains. We used 100 mg of bone powder, which was previously exposed to UV radiation for five minutes to reduce exogenous DNA contamination. Each sample was incubated in 2 mL of lysis buffer (0.5 M EDTA pH 8.00, 1 % N-Laurosylsarcosine sodium salt) and 50 µL proteinase K (20 mg/mL) for 24 h at 56 °C with gentle shaking. Samples were centrifuged for 5 min at 10,000 rpm.

DNA purification was performed using the E.Z.N.A Tissue DNA Kit (Omega Bio-tek, Norcross, GA, USA). A total of 1 mL of supernatant was recovered and divided into two 500 µL aliquots in sterile 1.5 mL microcentrifuge tubes. Subsequently, each aliquot was processed following the bind step suggested by the manufacturer’s protocol, but by adjusting the BL Buffer and 100% ethanol volume to 500 µL each. Two 700 µL filtrations were performed per column. Finally, DNA was eluted in 30 µL of Elution Buffer.

The DNA concentration and purity were measured using a NanoDrop spectrophotometer.

### PCR amplification

For the identification of cetacean species, the specific cetacean primers targeting cytochrome b gene (Lopez-Oceja et al. 2019), were employed. Forward primer L15601: 5’-TACGCAATCCTACGATCAATTCC-3’ and reverse primer H15748: 5’-GGTTGTCCTCCAATTCATGTTAG-3’.

Polymerase chain reaction (PCR) was carried out in a 20 µL volume containing 5 µL of 5X Taq Buffer, 1 µL of dNTPs (2.5 mM), 1.5 µL of each primer (10 µM), 1.5 µL of MgCl^2^ (25 mM), 0.5 µL of Taq DNA polymerase (Applied Biological Materials Inc. (abm)) (5 U/µL), 8 µL of nuclease-free water and 1 µL of DNA, except for the negative control, where 1 µL of nuclease-free water was added instead. The PCR protocol was as follows: initial denaturation for 2 min at 95 °C; 35 cycles of 30 s at 94 °C; 45 s at 54 °C; and 1 min at 72 °C; then a final extension for 7 min at 72 °C. PCR products were checked with 1.5% agarose gel electrophoresis. Finally, the PCR products were sent to Macrogen Inc. (Santiago, Chile) for sequencing in both directions.

### Molecular identification

The sequences obtained were edited and cleaned using the BioEdit program (v.7). Subsequently, the cleaned sequences were compared against the NCBI GenBank database using the BLASTn algorithm for species identification. Taxonomic assignment was evaluated based on Expect value (E-value), query coverage and percentage identity. As a complementary assessment of species assignment, a second BLASTn search was performed for each sequence excluding *Balaenoptera physalus* from the search set to evaluate similarity with alternative cetacean species.

To further assess taxonomic assignment, the recovered Cyt b sequences were aligned with reference sequences of *B. physalus* and representative mysticete species retrieved from GenBank using MUSCLE. Phylogenetic relationships were reconstructed in MEGA v.12 using the Maximum Likelihood (ML) method with 1,000 bootstrap replicates (Kumar et al. 2024). *Eubalaena japonica* was used as the outgroup.

## Results and Discussion

### DNA recovery and purity

DNA concentration in the present study ranged from 7.3 to 57.1 ng/µL (mean ± SD = 25.91 ± 13.91 ng/µL; Table 1), with a mean concentration higher than those reported in previous studies of cetacean bones. Ren et al. (2022) reported a concentration of 0.854 ng/µL from a Bryde’s whale vertebra (*Balaenoptera edeni*), sufficient to amplify and sequence the mitochondrial COI gene. Similarly, Dai et al. (2023) reported a range of 2.3 to 27.5 ng/µL (mean ± SE = 9.5 ± 6.8 ng/µL) from skull, scapula, vertebra and mandible fragments of cetaceans.

**Table 1.** DNA recovery and BLASTn molecular identification results for cetacean bone samples. ^a^ Skull and vertebra originated from the same individual. Dashes indicate samples with no success.

| Sample | Bone description | Bone powder (g) | DNA concentration (ng/μL) | Query coverage (%) | Identity (%) | E-value | First GenBank accession no. |
| --- | --- | --- | --- | --- | --- | --- | --- |
| 2200 | Skull | 0.119 | 28.0 | 98 | 91.26 | 1.0 x 10<sup>-59</sup> | KC572849.1 |
| 2198 | Mandible-fragment | 0.117 | 24.5 | 100 | 92.98 | 1.0 x 10<sup>-58</sup> | KC572854.1 |
| 2189 | Skull | 0.119 | 7.3 | 100 | 95.19 | 1.0 x 10<sup>-36</sup> | KC572849.1 |
| 2185 | Mandible | 0.125 | 21.6 | - | - | - | - |
| 2166 | Mandible-fragment | 0.110 | 35.1 | - | - | - | - |
| 2162 | Unknown | 0.111 | 18.5 | 100 | 95.19 | 1.0 x 10<sup>-36</sup> | KC572849.1 |
| 2152 | Vertebra | 0.117 | 57.1 | - | - | - | - |
| Huachipato<sup>a</sup> | Skull | 0.110 | 18.8 | 99 | 94.74 | 1.0 x 10<sup>-57</sup> | MT410921.1 |
|  | Vertebra | 0.115 | 22.3 | - | - | - | - |

DNA concentration variability could be attributed to differences in the density and composition of the sampled bone, as collagen and bone porosity are recognized determinants of DNA preservation in skeletal tissue. In cetaceans, compact bone is largely restricted to mandible and forelimb bones, whereas vertebrae and ribs are composed almost entirely of highly porous cancellous bone with only a thin cortical shell (Wysokowski et al. 2020). These structural difference may contribute to variation in DNA recovery, consistent with reports that bones with higher collagen content and a more compact microstructure tend to preserve amplifiable DNA more reliably (Sosa et al. 2013). In addition, DNA recovered from bone remains may be fragmented and present at low copy number, which may compromise subsequent molecular analyses (Pääbo et al. 2004).

DNA purity showed A260/280 ratios ranging from 1.46 to 2.11, with two values above the expected range of 1.8-2.0. A260/230 ratios ranged from 0.53 to 3.76, with two values above the optimal range of 1.8-2.2, potentially indicating the presence of co-extracted contaminants or residual extraction reagents (Desjardins and Conklin 2010; Żarczyńska et al. 2023). Sequencing failed for four samples (2185, 2166, 2152, and Huachipato vertebra), however, purity ratios alone did not clearly distinguish failed from successfully sequenced samples (Table S1). Notably, the Huachipato skull and vertebra showed comparable DNA concentrations (Table 1), although only the skull obtained showed sequencing success. Together, these observations indicate that neither DNA concentration nor purity ratios alone were predictive of sequencing success.

### PCR amplification and sequencing success

The Cyt b target was successfully amplified in all nine samples (100%), demonstrating that DNA recovered with the optimized protocol was amplifiable. Agarose gel electrophoresis (1.5%) revealed a single and well-defined band at approximately 500 bp for each sample (Fig. S1). Dai et al. (2023) reported amplification and sequencing success rates of 93.10% and 58.62% for the CR and CYB regions, respectively, from 29 cetacean bone samples using touch-down and nested PCR, whereas conventional PCR yielded considerably lower success rates (27.59% and 6.90%, respectively). Despite the 100% amplification rate obtained in the present study, only 55.6% of the samples (5 out of 9) were successfully sequenced, indicating that the presence of a visible PCR product did not guarantee sequencing success.

### Molecular identification

All successfully sequenced samples (n = 5) matched with *Balaenoptera physalus* reference sequences available in the NCBI GenBank database via BLASTn. Query coverage ranged from 98% to 100%, percentage identity from 91.26% to 95.19% and E-values from 1.0 × 10^-36^ to 1.0 × 10^-59^ (Table 1). Each bone sequence matched the species with more than 10 hits. In the complementary BLASTn analysis excluding *B. physalus*, second-best matches were inconsistent across samples, with no alternative species recurring with comparable support, further reinforcing *B. physalus* as the best match. Phylogenetic analysis further supported the BLASTn results, with all recovered sequences forming a well-supported clade closely associated with *B. physalus* reference sequences (100% bootstrap support) (Fig. S2). Together, these results provide additional functional validation that the optimized extraction protocol recovered mitochondrial DNA suitable for downstream molecular characterization.

## Acknowledgments

The authors thank the Servicio Nacional de Pesca y Acuicultura (SERNAPESCA), Blumar Fisheries and the Huachipato steel plant for providing the cetacean bone samples used in this study.

## Statements and Declarations

### Author contributions

DP: Formal Analysis, Investigation, Methodology, Visualization, Writing – original draft. SMR: Conceptualization, Funding Acquisition, Resources, Writing – review & editing. MG: Formal Analysis, Funding Acquisition, Project Administration, Resources, Supervision, Writing – review & editing.

### Data availability

Supporting data are available from the authors upon reasonable request.

### Ethics approval

No live animals were handled as part of this study. All bone remains were provided by local agencies.

## Supplementary Material

**Table S1.** DNA yield and purity metrics obtained from cetacean bone samples.

| Sample | Bone description | DNA |  |  |  |
| --- | --- | --- | --- | --- | --- |
|  |  | Concentration<br>(ng/μL) | A260 | A260/280 | A260/230 |
| 2200 | Skull | 28.0 | 0.559 | 1.86 | 1.57 |
| 2198 | Mandible-fragment | 24.5 | 0.489 | 2.11 | 3.25 |
| 2189 | Skull | 7.3 | 0.145 | 1.46 | 0.61 |
| 2185 | Mandible | 21.6 | 0.432 | 1.95 | 3.76 |
| 2166 | Mandible-fragment | 35.1 | 0.701 | 1.51 | 0.61 |
| 2162 | Unknown | 18.5 | 0.370 | 1.54 | 0.53 |
| 2152 | Vertebra | 57.1 | 1.143 | 1.51 | 0.58 |
| Huachipato | Skull | 18.8 | 0.376 | 2.08 | 2.17 |
|  | Vertebra | 22.3 | 0.447 | 1.93 | 1.61 |
| Mean | - | 25.91 | 0.518 | 1.77 | 1.63 |

**Fig. S1.**
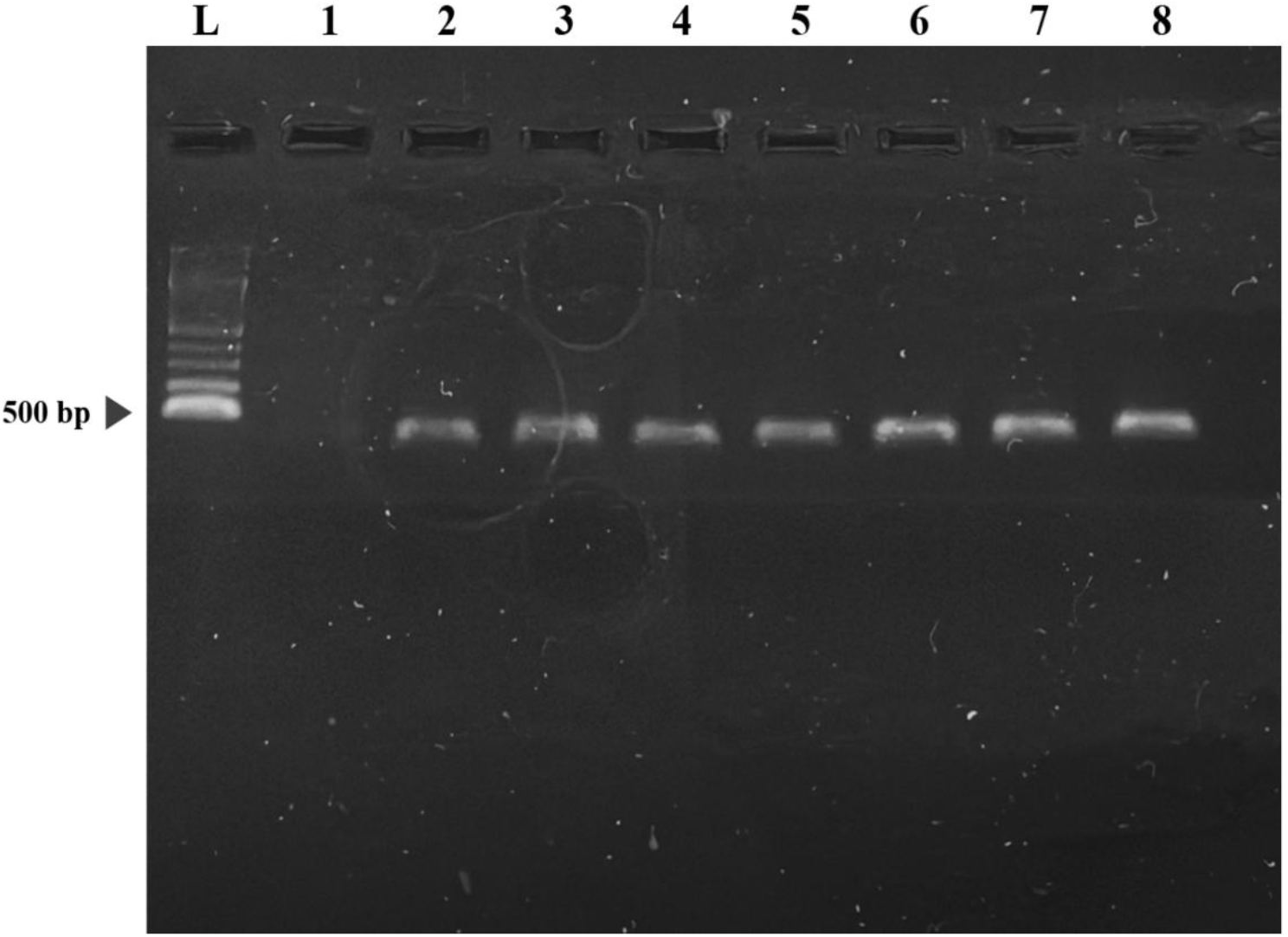
Agarose gel electrophoresis (1.5 %). Lane L: 100 bp Plus DNA Ladder (BIONEER). Lane 1: negative sample. Lanes 2-8: bone samples.

**Fig. S2.**
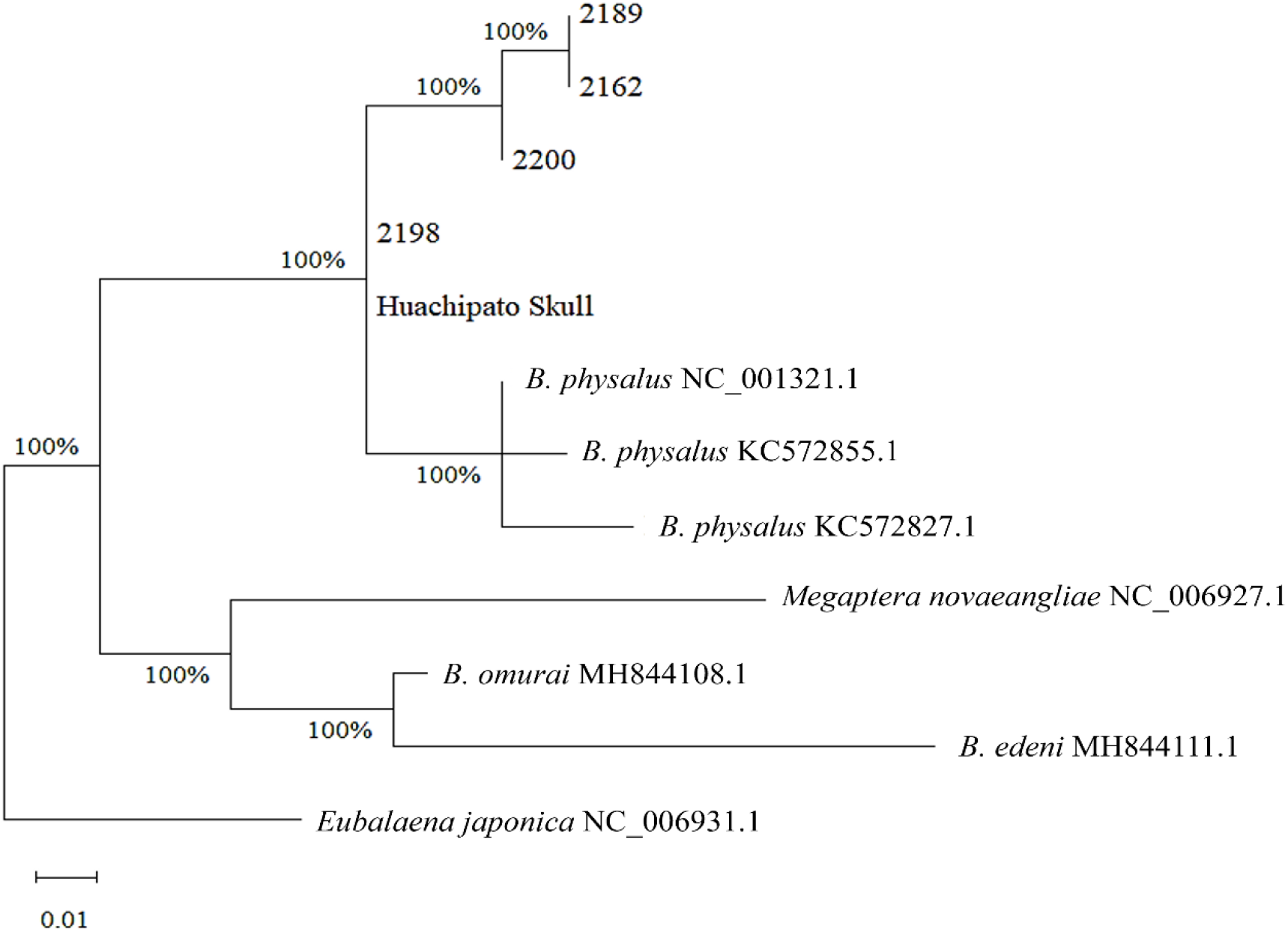
Maximum Likelihood phylogenetic tree based on mitochondrial Cyt b sequences recovered from cetacean bone samples and reference sequences retrieved from GenBank. Branch support was assessed using 1,000 bootstrap replicates. *Eubalaena japonica* was used as the outgroup. The scale bar represents 0.01 substitutions per site.

